# Vaeda computationally annotates doublets in single-cell RNA sequencing data

**DOI:** 10.1101/2022.04.15.488440

**Authors:** Hannah Schriever, Dennis Kostka

## Abstract

**Motivation:** Single-cell RNA sequencing (scRNA-seq) continues to expand our knowledge by facilitating the study of transcriptional heterogeneity at the level of single cells. Despite this technology’s utility and success in biomedical research, technical artifacts are present in scRNA-seq data. Doublets/multiplets are a type of artifact that occurs when two or more cells are tagged by the same barcode, and therefore they appear as a single cell. Because this introduces non-existent transcriptional profiles, doublets can bias and mislead downstream analysis. To address this limitation computational methods to annotate and remove doublets form scRNA-seq datasets are needed.

**Results:** We introduce vaeda, a new approach for computational annotation of doublets in scRNA-seq data. Vaeda integrates a variational auto-encoder and Positive-Unlabeled learning to produce doublet scores and binary doublet calls. We apply vaeda, along with seven existing doublet annotation methods, to sixteen benchmark datasets and find that vaeda performs competitively in terms of doublet scores and doublet calls. Notably, vaeda outperforms other python-based methods for doublet annotation. All together, vaeda is a robust and competitive method for scRNA-seq doublet annotation and may be of particular interest in the context of python-based workflows.

**Availability:** Vaeda is available at https://github.com/kostkalab/vaeda

**Contact:** @pitt.edu

## 1 Introduction

Single-cell RNA sequencing (scRNA-seq) continues to impact our understanding of diverse biomedical domains by providing high-resolution gene expression measurements at scale. Resulting datasets often comprise many thousands of cells or more; while they provide valuable insights, they are also limited by technical artifacts, like doublets/multiplets. Doublets or multiplets occur when two or more cells receive the same identifier during library construction and thus appear as one single cell. As such, doublets introduce nonexistent expression profiles, which can lead to incorrect interpretation of data in downstream analysis. While there are experimental techniques that identify and annotate doublets, these methods are currently not typically employed (reasons include an increase experimental burden and a decrease the cell yield), and they are not available for pre-existing datasets.

Therefore, computational methods to identify doublets are needed. Current methods that address this challenge include scDblFinder [1], doubletFinder [2], solo [3], Scrublet [4], and the cxds, bcds and hybrid methods of the scds approach [5]. Many of these methods share core concepts in their approach, for example generating artificial doublets from observed data, which then form the basis of deriving doublet scores for computational doublet detection. Here we propose vaeda, a new computational approach with a similar paradigm to existing methods; vaeda combines variational auto-encoders (VAEs, Kingma and Welling [6]) and Positive-Unlabeled Learning (PU-Learning, Liu et al. [7] and Mordelet and Vert [8]) to annotate doublets in scRNA-seq data. Similar to solo, vaeda uses a VAE to derive a low-dimensional representation of the input data. PU-Learning, on the other hand, is a learning framework designed for instances where there is a set of positively labeled examples and a set of unlabeled examples. In the context of doublet detection the unlabeled examples are input data, while the examples with labels are artificially generated doublets. Therefore, PU-Learning appears well-suited for doublet detection; however, to our knowledge, while PU-Learning has been used to identify cell-free droplets in scRNA-seq data by [9], it has not been used to identify doublets.

In the following we present the vaeda method in detail; we also apply it to 16 benchmark datasets with experimental doublet annotation [10] and show that vaeda can accurately annotate doublets in scRNA-seq data, performing well when compared to seven existing methods. We assess robustness of performance for all methods we study, and overall, we observe meaningful differences in how well the methods’ doublet scores correlate with experimental doublet annotations. Nevertheless, we also find that performance differences of binary doublet calls are less pronounced. This is of particular interest, because in practice, binary doublet calls are what enable researchers to remove artifacts and improve their data quality.

## 2 Methods

### 2.1 Vaeda method for doublet annotation

The vaeda method for doublet annotation consists of several steps summarized in **Fig**. 1. In the following we describe each step in more detail.

**Figure 1.**
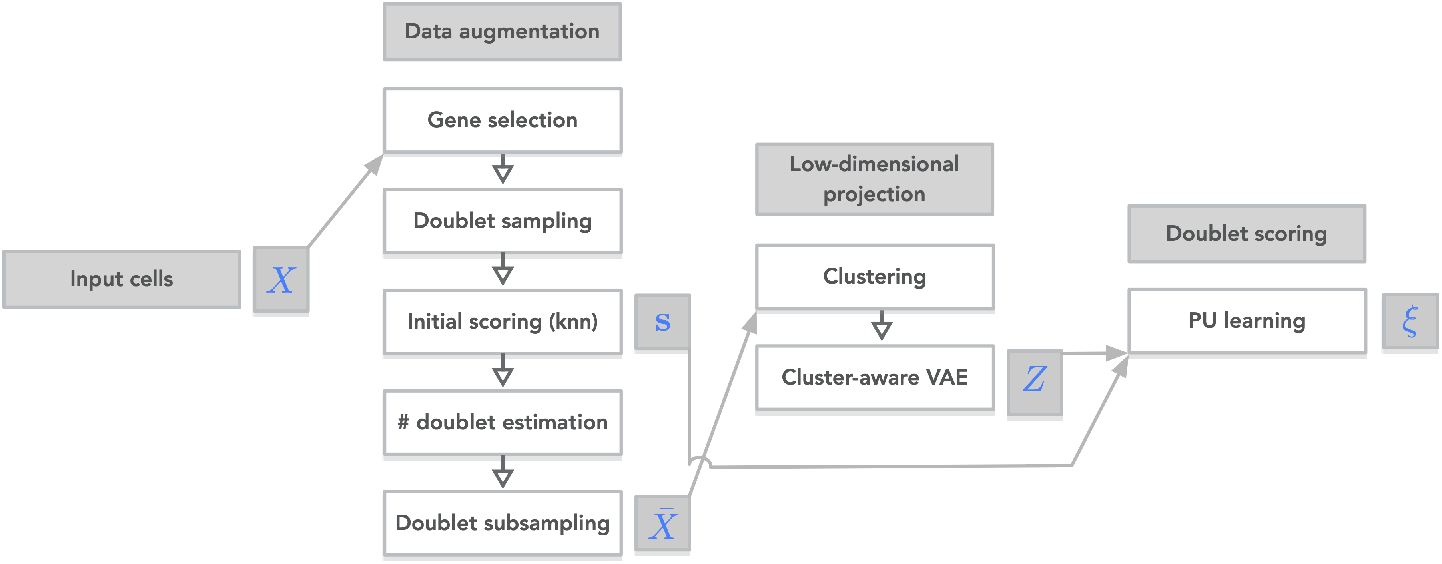
Summary of the vaeda method. Input cells *X* are subjected to data augmentation, where artificial doublets are simulated, a preliminary doublet score **s** is derived, and an augmented dataset 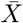 is created. Next, a low-dimensional representation *Z* of 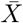 is derived, using a cluster-aware variational autoencoder. Finally, positive unlabeled learning is used to derive final doublet scores *ξ* for each input cell/barcode.

#### 2.1.1 Input data, doublet simulation, and gene selection

The input data for vaeda is a raw, un-normalized count matrix *X* with *n* rows (one per cell-barcode, encompassing singlets and doublets/multiplets) and *p* columns (one per gene). This data is then used to simulate artificial doublets as follows: An index pair (*i, j*) is sampled randomly and a doublet precursor 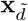 is created by adding the corresponding rows of 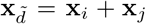. Next, to generate an artificial doublet **x**_*d*_, its library size *𝓁* (i.e., the number of counts) is determined as a random sample of all library sizes in *X* (i.e., the row sums of *X*) that are at least as large as the larger library size of **x**_*i*_ and **x**_*j*_. Then **x**_*d*_ is determined as 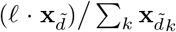. With this approach we create a count matrix *X*^*′*^ of *n* simulated doublets, and vaeda proceeds with the augmented matrix 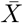 obtained by stacking *X* and *X*^*′*^ (and scaling and centering columns), and associated labels **y** indicating simulated doublets:

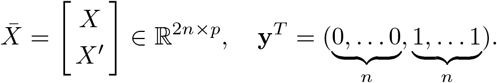

Finally, genes (i.e., columns) of 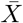 are selected by: (a) removing columns (i.e., genes) that had zero values for more than 99% of rows (i.e., cells) and then (b) focusing on the 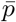 most variable columns (we choose 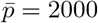), where variability is defined by variation. Overall this procedure yields an initial augmented expression matrix 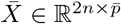.

#### 2.1.2 Adjusting the number of simulated doublets

Next, to adjust the number of simulated doublets in the augmented count matrix, vaeda uses the following approach. First, an initial estimate for the number of doublets present in the input data is derived. To that end, the augmented data 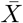 is projected on its first 30 principal components. Next, a *k* nearest neighbor (knn) graph is constructed, where we set 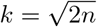. Then, for each cell the fraction of its nearest neighbors that are simulated doublets is calculated as a (preliminary) doublet score 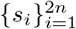. A score cutoff *c* is determined as the 25% quantile of simulated doublets’ scores. Finally, the number of doublets *n*_*d*_ in the original data is estimated as 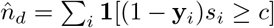, the number of input cells with scores equal or larger than the cutoff.

Second, 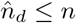 cells are randomly selected from simulated doublets by sampling without replacement, where the sampling probability for doublet *k* is given by 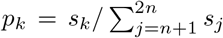. This results in 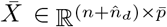 and **y** consisting of *n* zeros and 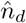 ones.

#### 2.1.3 Low-dimensional representation by cluster-aware variational auto-encoding

We derive a low-dimensional representation of the augmented data using a cluster-aware autoencoder; this representation will then in turn form the basis of doublet annotation.

##### Clustering

First, we group cells in the augmented dataset 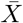 into clusters, using the Leiden algorithm [11]. Specifically, 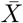 is scaled and projected onto its first 30 principal components. Next, for small datasets (1,000 cells or less), a neighborhood graph [12, 13] is computed followed by Leiden clustering. For larger datasets we pre-cluster the projected data using mini-batch k-means (similar to Hicks et al. [14], with *k* set to 10% of the number of cells) and generate meta-cells (i.e., cluster centers). Again, first 30 principal components of meta-cells are used to compute a neighborhood graph, followed by Leiden clustering. This step creates cell annotations 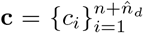 with *c*_*i*_ = *k* if cell *i* is assigned to cluster *k*.

##### Cluster-aware autoencoder

The cluster-aware autoencoder then takes 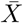 (log-transformed and scaled) and **c** as input and consists of an encoder network, a decoder network, and a cluster classifier. Let an input instance (i.e., cell and label) be denoted by (**x**, *c*).

The encoder network consists of an input layer (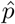 neurons), followed by a dense layer with 256 neurons, batch normalization and dropout (rate = 0.3) layers, and a tensorflow probability layer parameterizing a *d*-dimensional Normal distribution with diagonal covariance matrix (we use *d* = 5); *d* is the dimension of the latent space representation of the input data.

The decoder network consists of an input layer (*d* neurons), a dense layer of 256 neurons, followed batch normalization, dropout (rate = 0.3) layer, and a tensorflow probability layer parameterizing a *p*-dimensional Normal distribution with diagonal covariance matrix, modeling the input data, 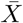.

The cluster classifier consists of an input layer (*d* neurons), batch normalization and a fully connected layer with the number neurons equal to the number of clusters present and with softmax activation, modeling each cell’s cluster assignment. Let the cluster classifier’s class assignments be denoted by *ζ*(**x**), and let *ϑ* denote the model’s trainable parameters.

The loss function of vaeda is defined as:

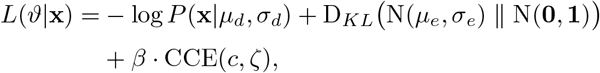

where the first two terms represent the loss function of the auto-encoder, and the third term is the classification loss of the cluster classifier (categorical cross entropy). Specifically, *µ*_*d*_(**x**, *ϑ*) and *σ*_*d*_(**x**, *ϑ*) represent output of the decoder’s final layer and are parameters of a Normal distribution modeling the input data, while *µ*_*e*_(**x**, *ϑ*) and *σ*_*e*_(**x**, *ϑ*) represent output of the encoder network’s final layer. Therefore, the first term denotes the negative log likelihood of the input data (=reconstruction error), while the second term is the Kullback-Leibler divergence between the (probabilistic) low-dimensional representation of the input and a standard Normal distribution of appropriate dimensions (=regularization). CCE is the categorical cross entropy between the input’s cluster label and the cluster classifier’s output *ζ*(**x**, *ϑ*), and *β* is a parameter adjusting the scale between autoencoder loss and classifier loss (we set *β* = 20, 000).

Vaeda is trained using the Adamax optimizer with default options. After the third epoch, the learning rate (0.001) decays at a rate of 0.75. Ten percent of input data is used as a validation set, and training stops if validation loss has not improved for 20 epochs. Moving forward, we use the auto-encoder’s low-dimensional representation of the input data 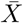 which we denote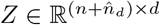.

#### 2.1.4 Doublet scoring by PU learning

To score cells as potential doublets we use a positive unlabeled learning approach, with the rationale that doublet labels on the input data are not observed (unlabeled), whereas we have positive labels on the simulated doublets in the augmented, reduced-dimensional data set (*Z*, **y**). Before we apply bagging PU learning, we augment (*Z*, **y**) by appending preliminary doublet scores **s** we used for the initial estimate of the number of doublets 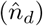, so that we finally use

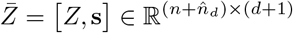

for doublet annotation. The PU bagging approach we use is summarized in Algorithm 1, which is adapted from the original publication [8]. Unlabeled examples *𝒰* are the first *n* rows of 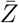, whereas the positive examples *𝒫* are the last 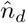rows of 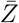. As classifier *f* (*x*) we use logistic regression, implemented as a neural neural network with an input layer (*d*+1 neurons), a batch normalization layer, and a dense output layer with sigmoid activation. We determine the number of epochs for training as follows: The network is trained on the first fold for 250 epochs; then the `KneeLocator` function of the `kneed` python module [15] is used to find an inflection point in the loss curve, which is then used to determine the number of training epochs for all folds. Scores 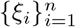 returned by PU bagging for the rows of 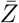 that represent input cells are the doublet scores reported by the vaeda method.

**Algorithm 1:**
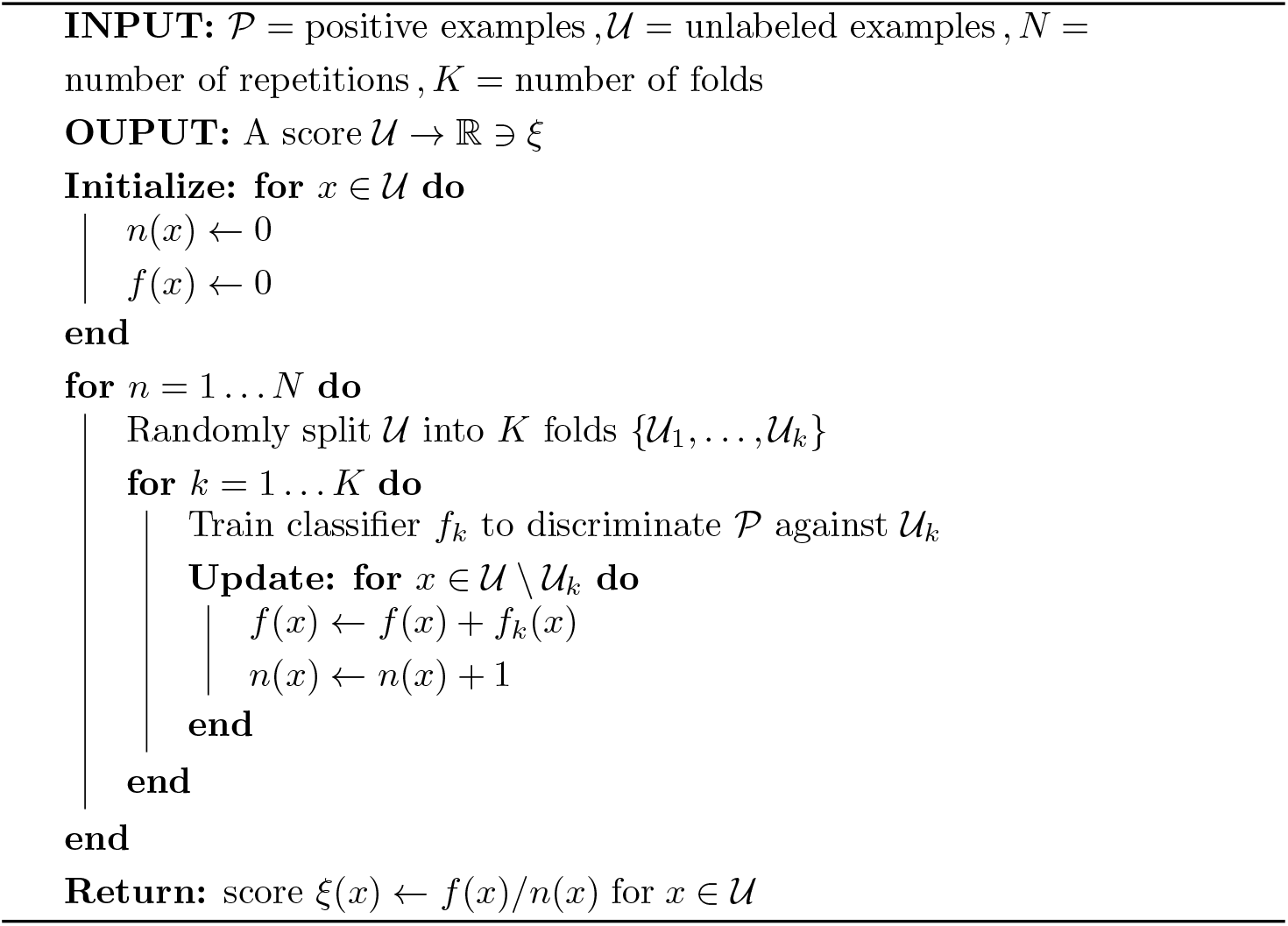
PU learning, adapted from Mordelet and Vert [8]

#### 2.1.5 Doublet Calling

In addition to doublet scores, vaeda also provides doublet calls as a binary prediction for whether a cell is a singlet or a doublet. Vaeda generates these calls by selecting a threshold *t*^*⋆*^ where all cells scoring above this threshold are called as doublets and all cells scoring below this threshold are called as singlets. The threshold is selected by the following minimization problem

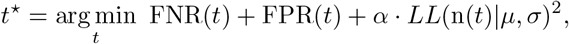

where FNR(*t*) is the fraction of simulated doublets called singlets at threshold *t* (*≈* false negative rate), FPR(*t*) is the fraction of input cells called doublets at threshold *t* (*≈* false positive rate), and n(*t*) is the number of input cells called doublets at threshold *t. LL*(n(*t*)|*µ, σ*) is the (Gaussian) log-likelihood of observing n(*t*) doublets given an expected number of doublets *µ* with standard deviation *σ*. We use *µ* = *n*^2^ *·* 10^*−*5^, motivated by the heuristic that *n ·* 10^*−*5^ is a good approximation for the fraction of doublets in a dataset (e.g., see [1]). Via a binomial model we arrive at a variance of *µ*(1 *− µ/n*) for the number of doublets, which determines the parameters of the log-likelihood term. We square the log-likelihood to flatten its minimum, and the parameter *α* controls a trade off between misclassification of simulated doublets and the number of expected doublets (we set *α* = *LL*(n(*t*_*max*_)|*µ, σ*)^*−*1^, where *t*_*max*_ is the largest possible threshold on the input data). We note that this approach is similar to that of Germain et al. [1]; however, the expected number of doublets is incorporated differently in our approach.

### 2.2 Benchmark data and application of existing methods

Scrublet [4], scDblFinder [1], doubletFinder [2], hybrid, bcds, and cxds [5] were run with default parameters using code from the R package DoubletCollection [16]. These methods also provide binary doublet calls in addition to continuous doublet scores, so we modified the package to access these calls for our doublet call analysis. Solo was run via the command line using parameters provided in the file `model_json` on solo’s github page (https://github.com/calico/solo). We use sixteen datasets from the benchmark study of Xi and Li [10], downloaded from Zenodo (https://zenodo.org/record/ 4062232#.X3YR9Hn0kuU).

The benchmarking datasets (pbmc-ch, cline-ch [17], mkidney-ch [3], hm-12k, hm-6k [18], pbmc-1A-dm, pbmc-1B-dm, pbmc-1C-dm, pbmc-2ctrl-dm, pbmc-2stim-dm, J293t-dm [19], pdx-MULTI, HMEC-orig-MULTI, HMEC-rep-MULTI, HEK-HMEC-MULTI, nuc-MULTI [20]) are summarized in Supplementary Table S1.

### 2.3 Analyses with down-sampled datasets

For several analyses we randomly down-sampled datasets ten times to contain a fraction of cells (we used 95%), while preserving the original singlet to doublet ratio. Each doublet annotation method was then run on each down-sampled dataset resulting in 10 annotation scores for each cell per method per dataset. Because of its longer running time solo was only run on five sub-samples instead of ten. To test for a difference in performance between two methods on a specific dataset we then used a Wilcoxon rank-sum test. To test for performance difference across all datasets we used a paired Wilcoxon ranksum test is used, where the pairing takes into account systematic differences between datasets. In cases of comparing solo to other methods, we only use the 5 repetitions where solo was applied.

To obtain a standard deviation of a performance measure averaged across datasets (e.g., **Fig**. 3 panel B) we estimated standard deviations for each dataset and then used standard error propagation.

**Figure 2.**
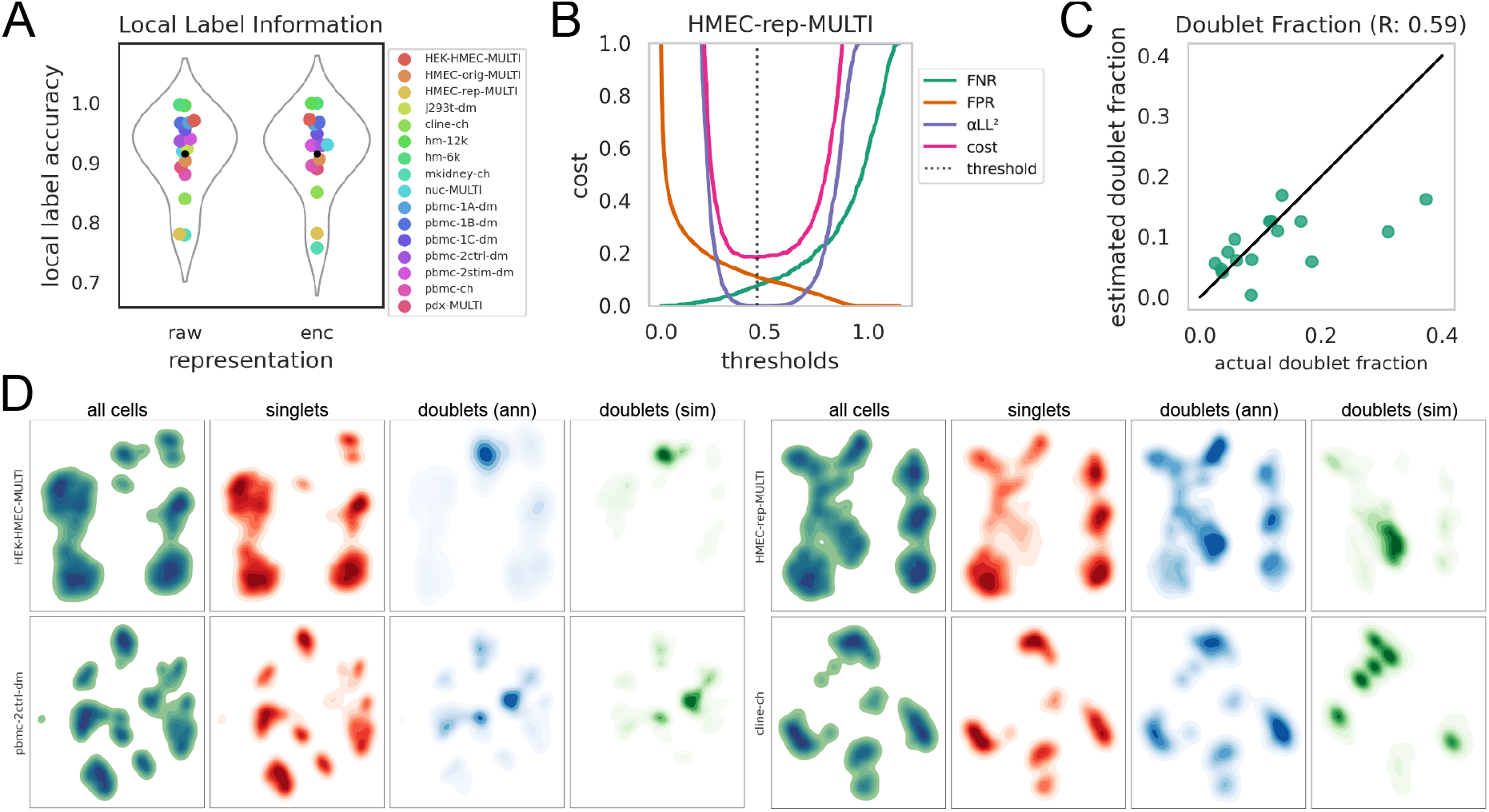
Vaeda’s latent representation preserves doublet annotation information and vaeda’s caller correlates with actual doublet fractions. Panel (A) shows the performance of a knn classifier predicting annotated doublets on the input data (left) and on vaeda’s latent representation (right). Panel (B) shows the cost function vaeda minimizes for doublet calling for the HMEC-rep-MULTI dataset. Panel (C) shows vaeda-estimated doublets on the x-axis and experimentally annotated doublets on the y-axis for 16 benchmark datasets. Panel (D) shows four datasets (rows) all cells, experimentally annotated singlets and doublets (columns two and three), as well as vaeda-simulated doublets (column four).

**Figure 3.**
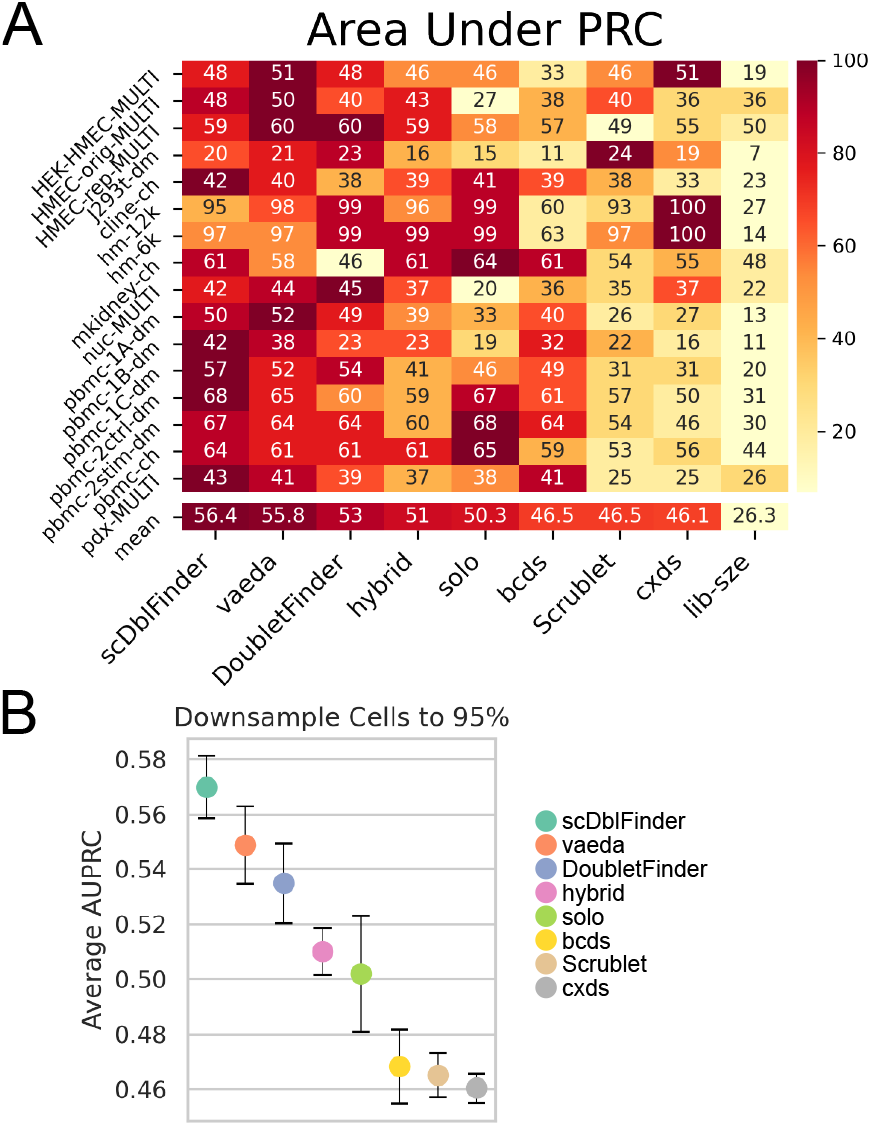
Vaeda performs well compared with other doublet annotations methods. Panel A shows average precision for doublet annotation methods (columns) across data sets (rows). The row labeled ‘mean’ denotes the average across datasets. Panel B shows average precision for repeatedly down-sampling 95% of cells across datasets for each method; error bars indicate three standard deviations.

### 2.4 Missed vs. captured doublets

We characterize and contrast experimentally annotated doublets we call “captured” and “missed” (**Fig**. 4). We define a captured doublet as a doublet that is classified as a doublet by at least four (out of eight) methods and a missed doublet as a doublet that is misclassified as a singlet by all of the methods. In this analysis we use for each method the *m* top-scoring doublets and as computationally annotated doublets, where *m* is the number of experimentally annotated doublets (i.e., we don’t use the methods’ doublet callers). For each doublet we then obtain the fraction of singlets amongst its k-nearest neighbors (k=5). **Fig**. 4 shows violin plots of these singlet fractions, stratified by missed vs. captured doubles. Circles represent averages for each dataset.

**Figure 4.**
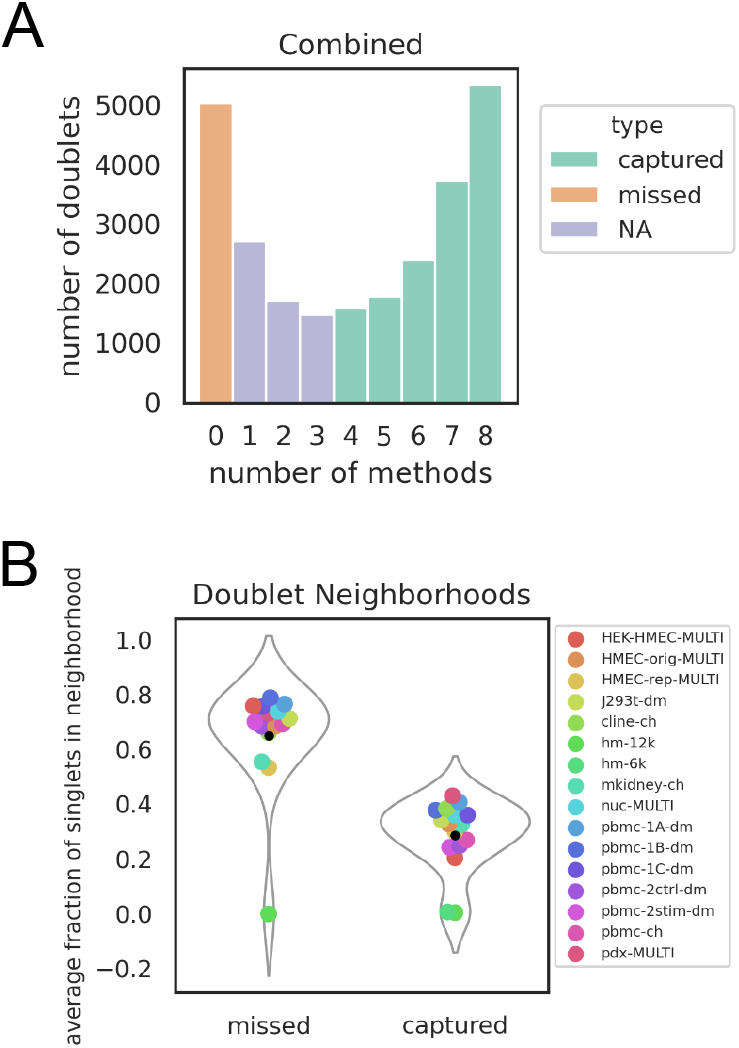
Missed and captured doublets. Panel (A) shows the number of experimentally annotated doublets, stratified by the number of methods that annotated them. Panel (B) shows density plots of the fraction of singlets in a doublet’s neighborhood, stratified by whether the doubled is consistently missed, or captured.

## 3 Results

### 3.1 Doublet detection with the vaeda method

Here we present the vaeda (Variational Auto-Encoder for Doublet Annotation) method for computational annotation of doublets in single cell RNA sequencing data. Vaeda reflects other doublet anno-tation methods ([1, 2, 5, 3, 4]), in that artificially generated doublets (we also call them “simulated” doublets) are used as a means to infer actual (or “real”) doublets in a dataset. Conditional on simulated doublets, this approach naturally leads to a statistical learning setup discriminating artificial doublets from input data, and classification scores are then used for doublet annotation. However, real doublets are present in the input data as well. With vaeda we explicitly account for this by viewing the learning task as a Positive-Unlabeled (PU) learning problem, where simulated data is viewed as a positive set (*P*), while input data is assumed unlabeled. This approach differs from standard classification, which implicitly assumes that no doublets are present in the input.

Briefly, the vaeda method works as follows. After variable gene selection and generation of artificial doublets, clustering is performed and a cluster-aware variational autoencoder (VAE) is used to learn a latent representation of input data and artificial doublets both. Learned latent projections, together with a knn-feature encapsulating the fraction of simulated doublets in each cell’s neighborhood, are then used as input for PU bagging [8] to derive doublet scores. Further on, vaeda can perform doublet calling. Similar to [1], vaeda balances a heuristically expected number of doublets with false positive and false negative doublet calls, as quantified by the number of misclassified simulated doublets and input cells, respectively. See Methods section for details and **Fig**. 1 for an overview of vaeda. In the following we assess vaeda on 16 benchmark datasets [10] where doublets have been experimentally annotated.

#### 3.1.1 Ablation analysis of the vaeda method

In order to evaluate the relative importance of vaeda’s different components we performed ablation analyses. Specifically, we assessed the following components of our method: 1) inclusion of the fraction of (simulated) doublets in a cell’s neighborhood into the learning problem, vs. not including them; 2) classification algorithm: knn classifier vs. logistic-regression classifier; 3) excluding simulated doublets that might be homotypic, ie doublets simulated by combining two cells from the same cluster; 4) type of low-dimensional representation: pca vs. variational auto-encoder vs. cluster-aware variational autoencoder; 5) PU learning vs. regular classification. We measured performance for every combination of these components, and results are summarized in Supplemental **Fig**. S1.

We find that a combination of a cluster-aware autoencoder, homotypic doublet exclusion, PU learning with a logistic regression type classifier, and including the neighborhood doublet fraction as a feature yielded the best results (average Area under the Precision Recall Curve: 55.8%, see **Fig**. 3). We also find that combinations including the neighborhood fraction of doublets and a logistic regression type classifier perform best, while methods without this feature using a logistic regression classifier perform worst. Combinations with a knn-type classifier perform in-between. Focusing on twelve high-performing combinations (i.e., with neighborhood doublet fraction and logistic regression classification), we find that combinations including PU learning perform better than those using regular classification (four of the top-six and two of the top-three combinations use PU learning). Results with regards to the low-dimensional embedding are a bit less clear; however, two of the top-three combinations use a cluster-aware vae (one uses PCA), while only one of the worst-performing combinations uses this approach for dimension reduction (the other two methods used are PCA and a regular auto-encoder). Overall this analysis motivates our design of the vaeda method. While biggest effects came from the neighborhood doublet fraction and classifier type, we note that PU learning and the cluster-aware autoencoder increased average performance from 54.5% to 55.8%, a mild but noticeable improvement.

#### 3.1.2 Vaeda’s latent representation preserves doublet annotation

Vaeda learns a latent space representation of the data by training a cluster-aware VAE. We quantitatively and qualitatively assessed how well the latent representation preserves experimental doublet annotation. First, to quantitatively assess whether this embedding preserves local label (=doublet) information, we followed the approach of [21] and use a knn (k=5) classifier trained using experimental doublet annotations to predict cell labels in both the input space and in the latent space. We find that label accuracy in vaeda’s latent space is 91.598% (averaged over datasets), approximately the same as in the input space (91.586%) (see **Fig**. 2). This indicates that vaeda’s latent representation accurately reflects local cell label information.

Next, to qualitatively assess vaeda’s simulation of artificial doublets, we visualized simulated and experimentally annotated doublets using vaeda’s latent representation. Plots for all benchmark datasets are shown in **Fig**. S2; **Fig**. Overall, we observe good agreement between experimentally-annotated and simulated doublets. However, for some datasets (HMEC-rep-MULTI, cline-ch, mkidney-ch, pbmc-1B, and pbmc-1C) there exist doublet populations that are annotated but not covered well by simulated doublets. We also note that regions with high densities of singlets are typically distinct from high-density simulated doublet regions. Two exceptions are hm-6k and hm-12k, which both have small groups of simulated doublets overlapping annotated singlets. We note that experimental doublet annotations for these data do not consider homotypic doublets (i.e., doublets that occur when two cells of the same cell type combine). **Fig**. 3D shows two examples of good real-simulated doublet overlap on the left, and HMEC-rep-MULTI and cline-ch on the right.

#### 3.1.3 Vaeda’s doublet scores and doublet calling reflect experimental annotations

Doublet scores produced by the vaeda method correlate well with experimental doublet annotation. Briefly, across 16 benchmark datasets, vaeda achieves an average area under the precision recall curve (AUPRC) of 55.8%, with with AUPRCs for individual datasets reaching 97% and 95% (**Fig**. 3). More details about vaeda’s doublet score performance and a comparison with other methods are presented in the next section.

In addition to calculating doublet scores for each cell, vaeda can also classify cells as either a doublet or a singlet as described in Section 2.1.5. Briefly, vaeda’s doublet caller simultaneously minimizes false positive rate, false negative rate, and the squared log likelihood of the predicted number of doublets given an expected number of doublets. The expected number of doublets is derived from the observation that, across experiments, the probability of a cell being a doublet scales linearly with the total number of cells, also see Section 2.1.5. **Fig**. 2 shows vaeda’s doublet calling results, compared to the number of experimentally annotated doublets. The correlation between annotated and estimated doublet rates (*r*^2^ = 0.59) is a noticeable improvement over using purely the expected number discussed above (*r*^2^ = 0.52, Supplementary **Fig**. S4). Further on, performance on datasets with reasonable doublet rates (less than 20%) appears more accurate, while the doublet fraction in two data sets with high doublet density (37.3% and 31.0%) is underestimated.

In summary, these results demonstrate that the vaeda method is a reasonably-designed method that generates accurate doublet scores for scRNA-seq data; it is able to perform meaningful classification into singlet vs. doublet cells, outperforming a reasonable baseline based on cell-numbers alone. Next, we compared vaeda with other computational methods for doublet annotation.

### 3.2 Comparison with other Doublet Annotation Methods

To compare vaeda to other methods, we used 16 benchmarking datasets that have previously been used to assess doublet annotation performance by [10]. We proceed by comparing vaeda with scDblFinder, doubletFinder, hybrid, bcds, cxds, solo, and scrublet, while library size was included as a baseline. We used average precision (area under the precision recall curve, AUPRC) as the main performance metric, because the datasets are imbalanced (typically the fraction of doublets is low, see Table S1). Results are summarized in **Fig**. 3 (panel A).

We find that vaeda outperforms all competing methods on four datasets; this is slightly worse than scDblFinder (five), but better than doubletFinder (two), Solo (three), cxds (three) and Scrublet (one). Hybrid and bcds did not outperform all other methods on any dataset. In terms of average performance, vaeda’s AUPRC (averaged across datasets) is 55.8%, second compared with 56.4% for scDblFinder (the best method, on average) and 53% for doubletFinder (the number three method), see **Fig**. 3, panel A.

#### 3.2.1 Vaeda performs competitively with existing methods

Performance results presented in the previous section were computed using all cells in each benchmark dataset. To study if the performance differences we observed were robust and meaningful, we sub-sampled cells in each benchmark dataset repeatedly and observed the resulting empirical distribution of performance metrics. In **Fig**. 3, panel B shows results using repeated (ten times) 95% down-sampling in terms of average AUPRC. Examining average performance, we find that vaeda outperforms all other methods, except scDblFinder and doubletFinder. We observe a noticeable decrease in average performance between the top three methods (scDblFinder, vaeda, doubletFinder) and other methods. Similarly, bcds, Scrublet and cxds on average perform worse than the rest. We also quantitatively assessed performance differences using Wilcoxon rank-sum tests for pairs of methods on the 95% down-sampled datasets, aggregating across all 16 datasets (see Methods, Section 2.3); we find that (using continuous doublet scores and AUPRC) scDblFinder outperforms vaeda and vaeda outperforms doubletFinder (Table 1).

**Table 1:**
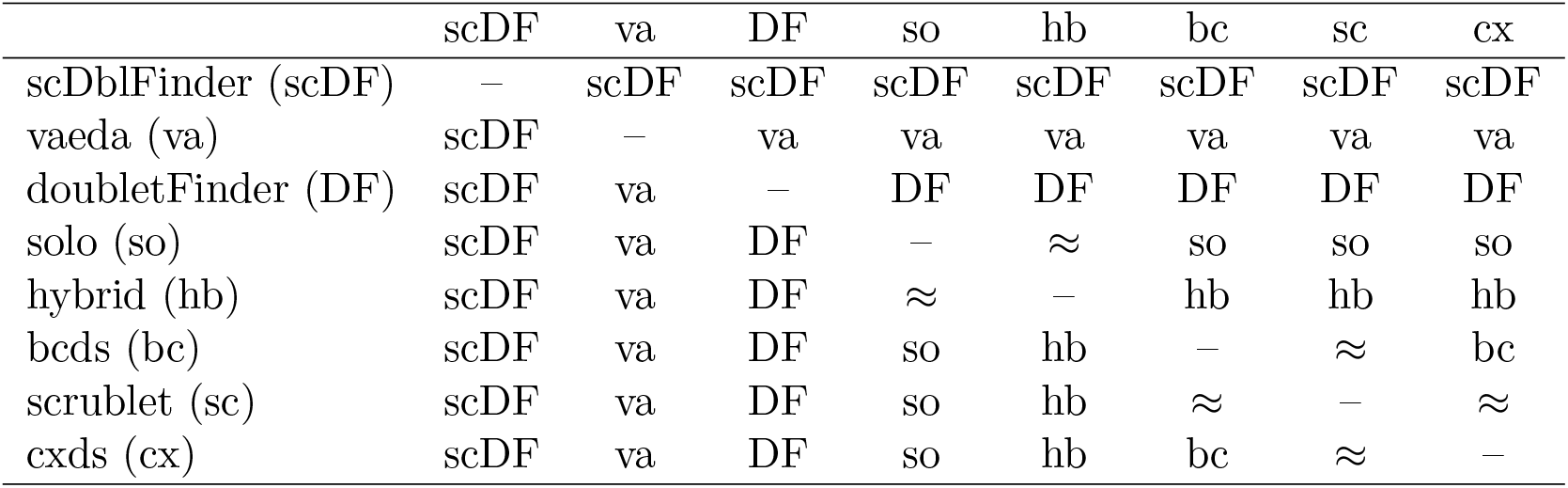
Comparison of methods aggregated across benchmark datasets. Paired Wilcoxon rank-sum tests were used identify significant (*p ≤* 0.05) performance differences between pairs of methods using 10 95% down-samplings of each dataset. The method with the better performance is indicated. *≈* implies *p >* 0.05.

We also compared all pairs of doublet detection methods stratified by dataset, using paired Wilcoxon rank-sum tests to decide “wins” (*p ≤* 0.05, higher performance), “ties” (*p >* 0.05), and “losses” (*p ≤* 0.05, lower performance). Panel C in **Fig**. S7 shows the results. For each method, there are seven competitor methods and 16 datasets yielding 7 *×* 16 = 112 comparisons. Only scDblFinder, vaeda, and doubletFinder are able to win more than half of their comparisons (83, 67 and 64, respectively). For the top-three methods (scDblFinder, vaeda, doubletFinder) we also provide pairwise comparisons, stratified by dataset, in Supplemental Table S2.

#### 3.2.2 vaeda performs competitively at doublet calling

In practice, doublet annotation methods are used to filter putative/annotated doublets from scRNA-seq datasets. In addition to doublet scores, vaeda is equipped with a doublet caller that provides a binary label for this purpose. Here we compare vaeda’s doublet calling with other methods and find that it performs competitively. Specifically, we applied vaeda along with other methods to the 16 benchmark datasets and recorded doublet calls. We then calculated the f1 score, mcc, precision, recall, and accuracy for the calls and averaged across datasets (Table 2, Supplementary **Fig**. S8). We see that vaeda performs well overall, with the second highest f1-score; compared to scDblFinder, vaeda appears to trade off recall (52.5% vs. 56.9%) to gain precision (59% vs. 53.7%), but coming out a little behind overall. Interestingly doubletFinder does not perform well, calling too few doublets overall. We also observe that there is noticeable spread in performance between different data sets (Supplemental **Fig**. S8).

**Table 2:** Doublet calling across 16 benchmark datasets (averaged performance).

|  | f1 | mcc | precision | recall | accuracy |
| --- | --- | --- | --- | --- | --- |
| scDblFinder | 51.4 | 47.7 | 53.7 | 56.8 | 89.3 |
| vaeda | 49.6 | 46.8 | 59.0 | 52.5 | 89.2 |
| hybrid | 47.9 | 4.30 | 48.7 | 53.3 | 87.8 |
| solo | 47.6 | 43.3 | 48.8 | 54.8 | 88.0 |
| bcds | 45.3 | 40.9 | 49.1 | 48.6 | 88.0 |
| cxds | 39.6 | 33.7 | 39.3 | 48.3 | 85.1 |
| Scrublet | 35.2 | 37.5 | 80.4 | 27.8 | 89.3 |
| lib-size | 27.5 | 20.6 | 29.5 | 30.7 | 84.5 |
| doubletFinder | 0.7 | 4.0 | 87.5 | 0.3 | 87.8 |

To investigate whether calling doublets is better than using the expected number of doublets, based on the heuristic *n*^2^ *·* 10^*−*5^ (see Methods, Section 2.1.5) to select a cutoff, we recalculated performance metrics for all methods with this approach. Supplemental Table S3 shows the results, and Supplemental Table S4 shows the difference in performance. For most methods performance differences between this approach and method-specific determination of the number of doublets are small; exceptions are Scrublet and doubletFinder, with both of them losing precision and gaining recall as more doublets are called.

To test for significant differences in performance in doublet calling we used Wilcoxon rank-sum tests, like before. Table 3 shows results comparing scDblFinder, vaeda and doubletFinder for f1 score, mcc, precision, recall and accuracy. We find that vaeda performs comparable to scDblFinder in terms of mcc, precision and accuracy, and slightly worse in terms of f1 score and recall.

**Table 3:** Comparison of doublet calls averaged across benchmark datasets. Paired Wilcoxon rank-sum tests were used identify significant (*p ≤* 0.05) performance differences between methods’ doublet calls. *≈* means *p >* 0.05.

|  | f1 | mcc | precision | recall | accuracy |
| --- | --- | --- | --- | --- | --- |
| vaeda-scDblFinder | scDF | ≈ | ≈ | scDF | ≈ |
| vaeda-doubletFinder | vaeda | vaeda | ≈ | vaeda | ≈ |
| scDblFinder-doubletFinder | scDF | scDF | DF | scDF | ≈ |

#### 3.2.3 Consistently misclassified doublets may be homotypic

In a majority of benchmarking datasets, there is a large subset of annotated doublets that are misclassified by every method (**Fig**. S9, S10). We suspect that these consistently misclassified doublets, or “missed” doublets, are composed primarily of homotypic doublets (i.e., doublets that are composed of two cells of the same type). Support for this hypothesis comes from the fact that the only two datasets that do not have a substantial amount of (consistently) missed doublets are the hm-6k and hm-12k datasets, where homotypic doublets are not annotated. In order to further test this idea, we measured the mixing between singlets, captured doublets, and missed doublets; in this analysis captured doublets are annotated by at least four methods, and missed doublets were not annotated by any of the eight methods we applied. Mixing was measured as the average fraction of singlets in the neighborhoods of the missed/captured doublets as described in Section 2.4. We found that missed doublets have higher mixing with singlets than captured doublets, providing support to the idea that the missed doublets are homotypic (i.e., near singlets presumably of the same type), see **Fig**. 4.

## 4 Discussion

Here we present vaeda, a new tool for computational doublet detection. The vaeda method uses a similar paradigm to other approaches, where doublets scores are produced using artificially generated doublets (e.g., Bais and Kostka [5], Germain et al. [1], and others). While auto-encoders have been used in the context of doublet detection before [3], vaeda is unique in that it combines a cluster-aware variational auto-encoder with a PU-Learning approach. We carefully studied the effect of both of these concepts and showed that, even though other design choices have greater impact, they do enable a noticeable increase in performance.

We assessed vaeda on 16 benchmark datasets, and find that, overall, vaeda produces accurate doublet scores and is able to derive binary doublet predictions that reflect experimental annotations well. Further on, we find that vaeda’s latent representation is helpful in determining datasets where simulated doublets agree well with experimental annotations, and cases where simulated and experimentally annotated doublets show more pronounced differences. We illustrate this with four examples in **Fig**. 2 D, with HMEC-rep-MULIT and cline-ch as examples where the distributions simulated and annotated doublets show differences. We note that vaeda, as well as other methods (see **Fig**. 3), do not perform particularly well on these data, indicating that improving our ability to simulate doublets might be a promising approach to improve method performance.

In terms of assessing method performance, we note that ultimately the metrics most relevant in practice are those for binary predictions, rather than rank-based metrics like area under the ROC curve or average precision (i.e., area under the precision recall curve). Such metrics are reported in Table 2, and we see that vaeda outperforms doubletFinder and performs comparable so scDblFinder in terms of mcc, precision and accuracy; in terms of f1 score and recall scDblFinder performs a bit better than vaeda. Therefore we conclude that vaeda performs comparable to other state-of-the-art methods (scDblFinder and doubletFinder), but note that scDblFinder has slightly better average performance in terms of f1 score and recall. We highlight, however, that the only other native python method is scrublet, which performs significantly worse than vaeda.

In summary, we have shown that vaeda is a state-of-the-art method for computationally annotating doublets in scRNA-seq data, and that it is the top choice for python workflows. It also provides a low-dimensional data representation that can complement other approaches and be useful for data analysis and visualization. It therefore is a useful tool for single cell RNA sequencing data analysis.

## Acknowledgements

HS and DK would like to thank the Kostka and Chikina labs for feedback and discussion.

