## Supplemental Information for "Vaeda computationally annotates doublets in single-cell RNA sequencing data"

For the manuscript

#### S1 Supplemental Figures

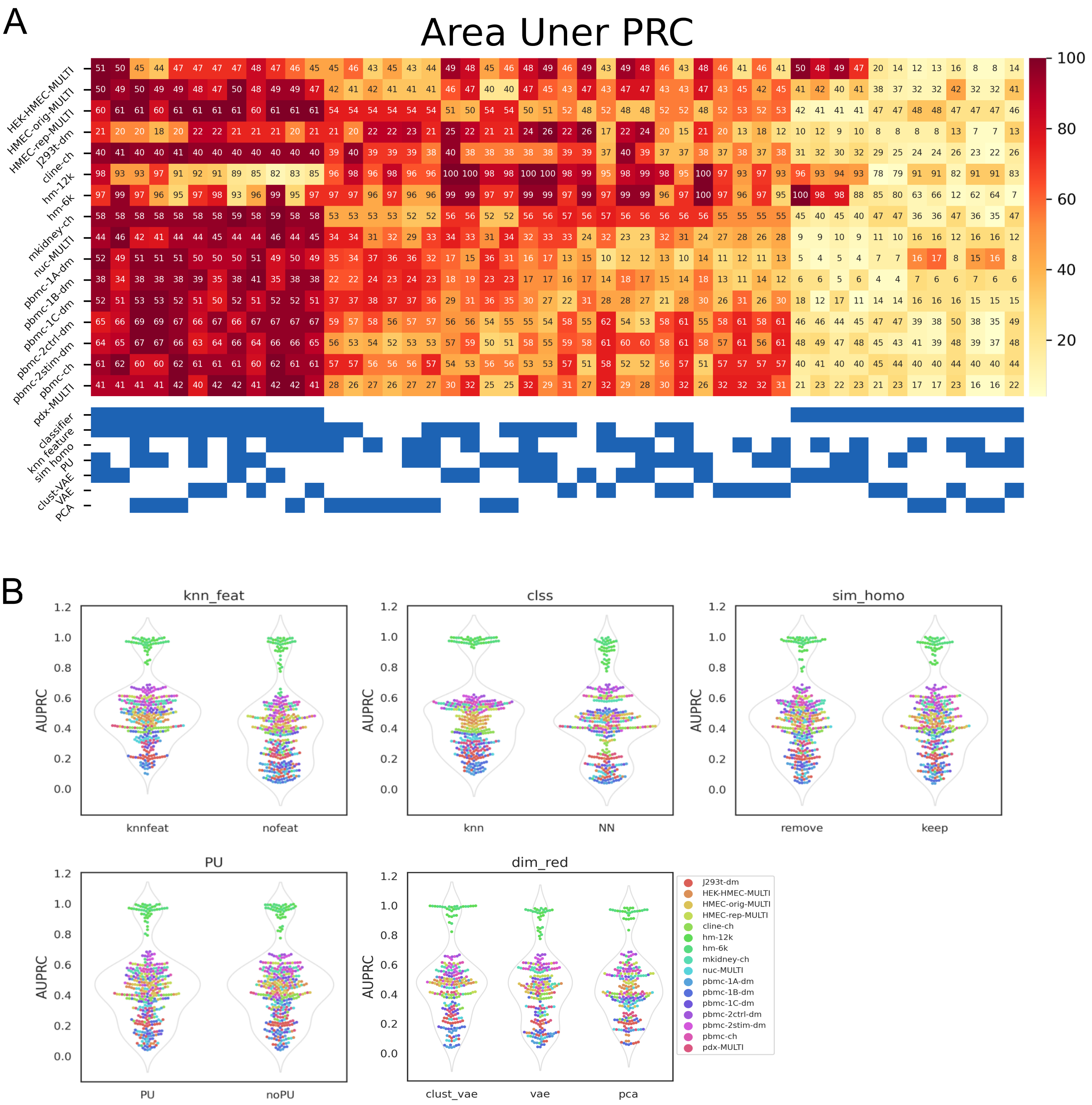


Supplemental Figure S1: ***Ablation Analysis shows the importance of combining knn and neural network classifier.*** (A) Heat map showing area under the precision recall curve for every permutation on the components of vaeda (columns) on each benchmarking dataset (rows). Colors are normalized by row and correspond to performance with burgundy being the best and pale yellow being the worst. Bottom chart depicts which components are used. The key is as follows. classifier: blue=neural network classifier, white=knn classifier; knn_feature: blue=append to low dimensional representation, white=no knn_feature; sim_homo: blue=include simulated homotypic doublets, white=remove simulated homotypic doublets; PU: blue=PU loop implemented, white=PU not used. Then for clust_vae, vae, and PCA blue indicates that methods was used for dimensionality reduction and white indicates it was not. (B) Violin plots showing AUPRC for every permutation on the components of vaeda, with each plot focusing on the significance of one component.

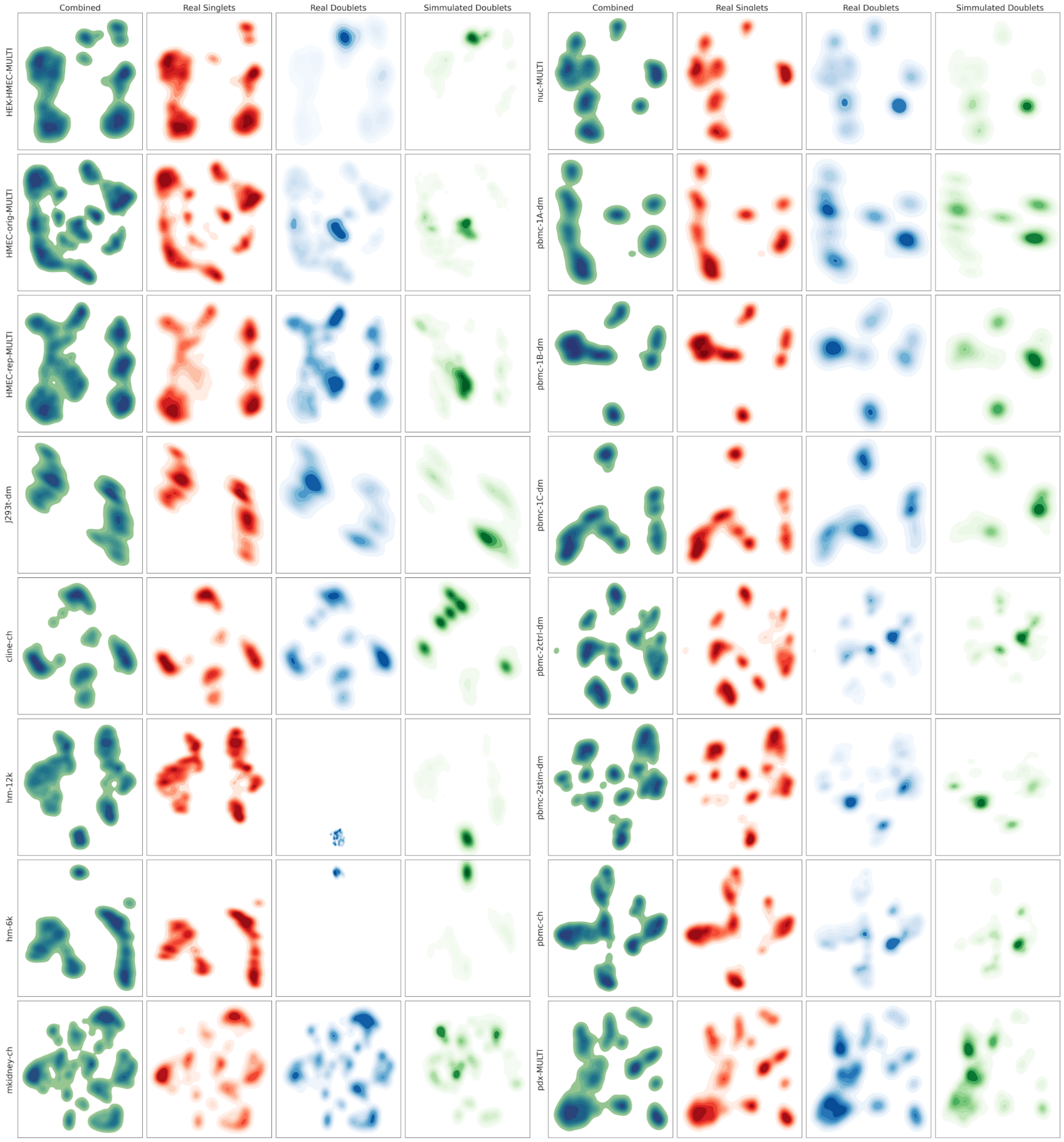


Supplemental Figure S2: ***Simulated doubles largely agree with experimentally annotated doublets.*** Shown are two-dimensional projections of vaeda's latent representation of cells. Rows are different benchmark datasets. The first column in each panel are all cells in the dataset; the second and third columns show experimentally annotated singlets and doublets, respectively; the third column shows vaeda's simulated doublets.

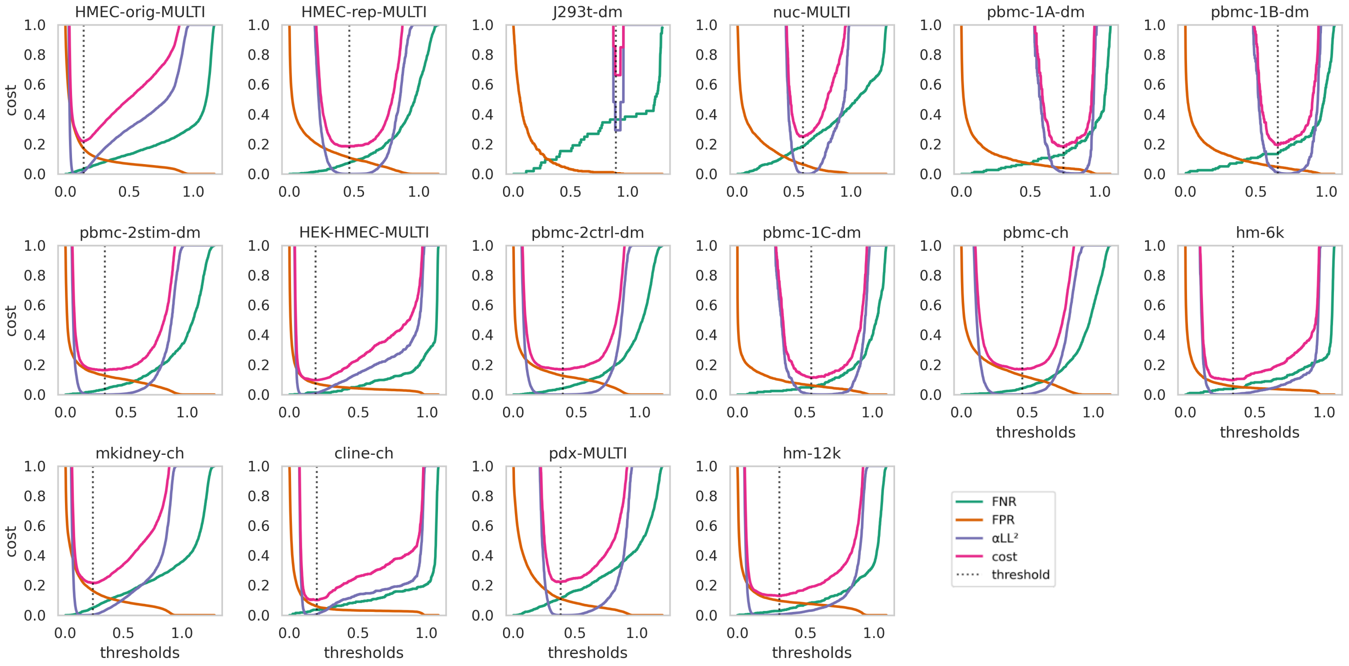


Supplemental Figure S3: ***Cost functions used for thresholding during doublet calling.*** Depicted above are the cost functions, along with the components of the cost functions for each dataset. FNR=false negative rate; FPR=false positive rate; $LL^2$=likelyhood squared; cost=FNR+FPR+$LL^2$; threshold=minimum of the cost function.

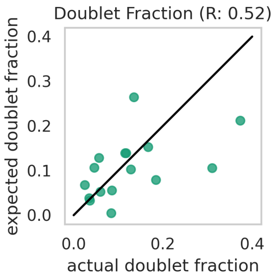


Supplemental Figure S4: ***Expected number of doublets vs actual number of doublets.*** Each point represents one of the benchmarking datasets. Expected doublet fraction correlates with actual doublet fraction with a correlation coefficient of 0.52.

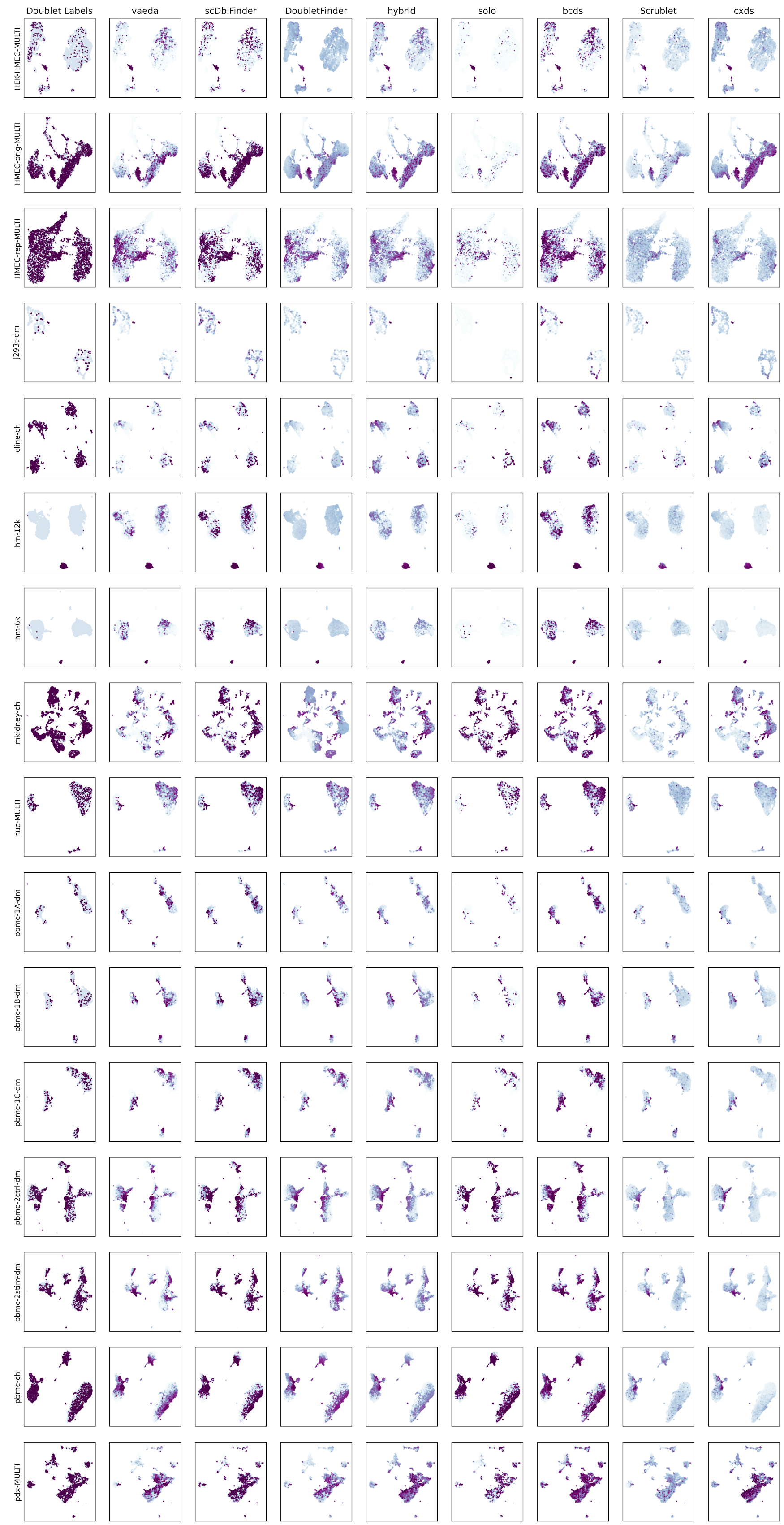


Supplemental Figure S5: ***Score distributions from each method projected onto UMAP representations of each benchmarking dataset.*** Depicted above are UMAP projections on the top 30 Principal Components of each dataset. Rows are datasets, the first column is the experimental doublet annotations, and the remaining columns each are a method. In the first column, purple=doublet; pale blue=singlet. In the remaining columns, dark purple represents higher doublet scores while pale blue represents lower scores.

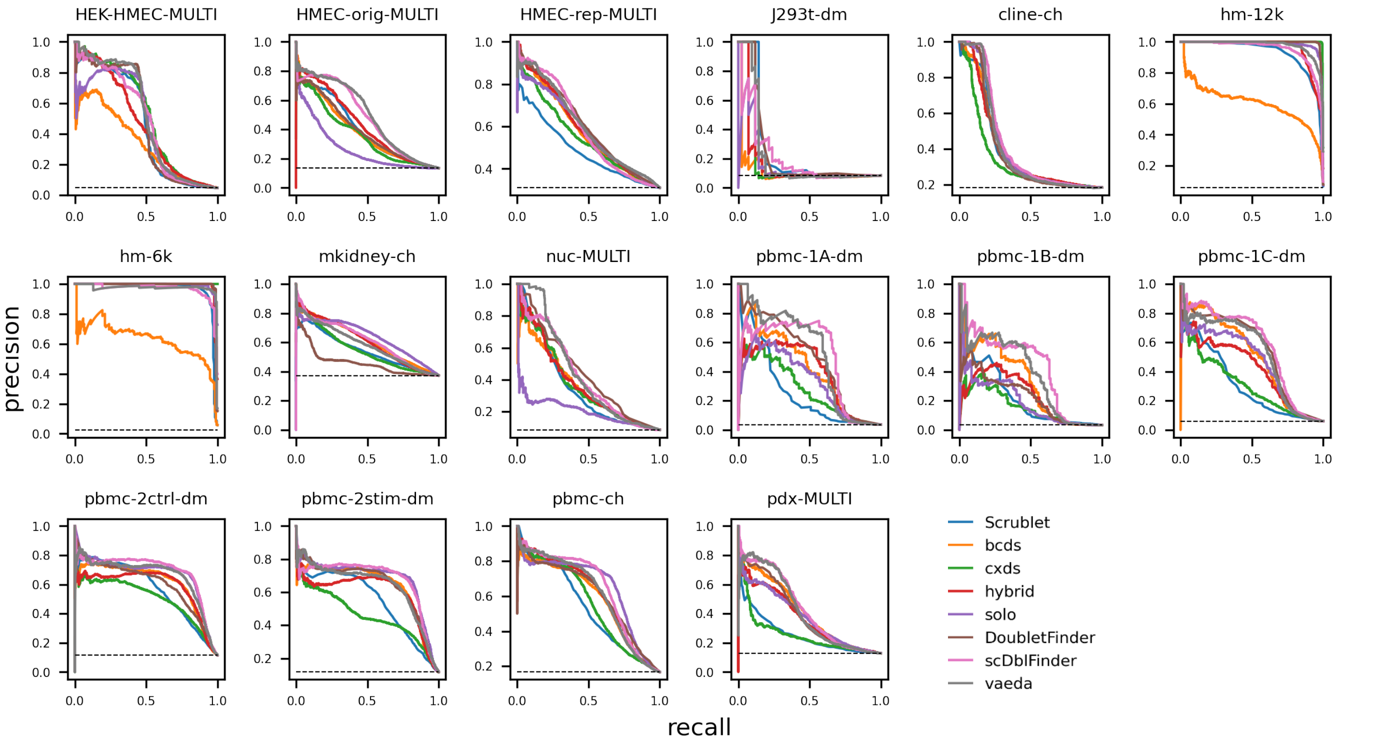


Supplemental Figure S6: ***Precision Recall Curves for each method.*** Shown above are the Precision Recall Curves for each method applied to each dataset.

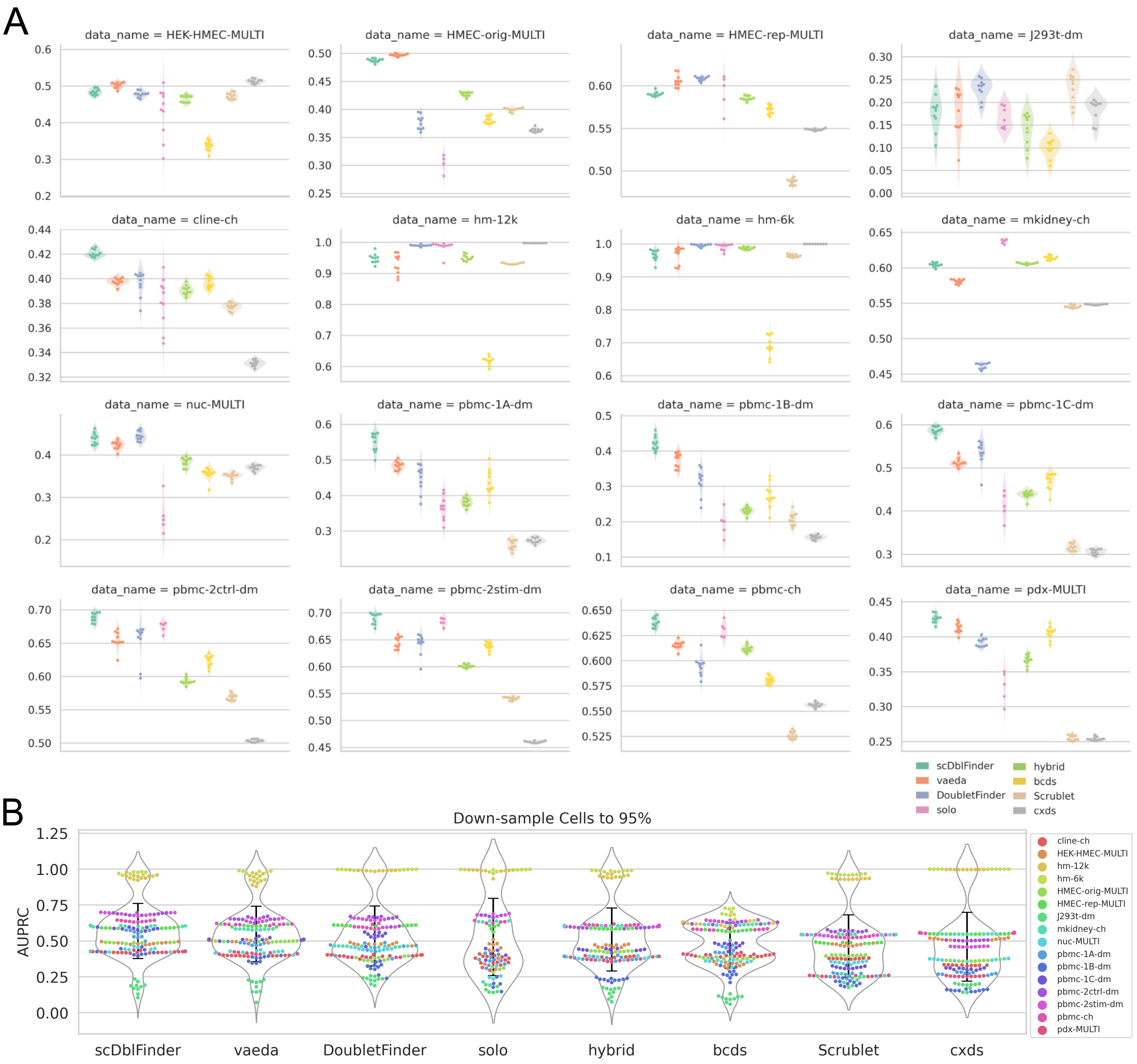


C

| method | wins | ties | losses |
|---|---|---|---|
| **scDblFinder** | 83 | 13 | 16 |
| **vaeda** | 67 | 17 | 28 |
| **DoubletFinder** | 64 | 16 | 32 |
| **solo** | 46 | 23 | 43 |
| **hybrid** | 43 | 13 | 56 |
| **bcds** | 41 | 8 | 63 |
| **Scrublet** | 20 | 10 | 82 |
| **cxds** | 32 | 4 | 76 |

Supplemental Figure S7: ***Area under the Precision Recall Curve for methods applied to down-samplings of each dataset.*** (A) Violin plots depicting area under the Precision Recall Curve for each down-sampling across methods and datasets. (B) Violin plots of AUPRC for each method on 95% down-sampled datasets. (C) Table showing all pairwise comparisons of doublet detection methods stratified by dataset, using paired Wilcoxon rank-sum tests to decide "wins" ($p \leq 0.05$, higher performance), "ties" ($p > 0.05$), and "losses" ($p \leq 0.05$, lower performance)

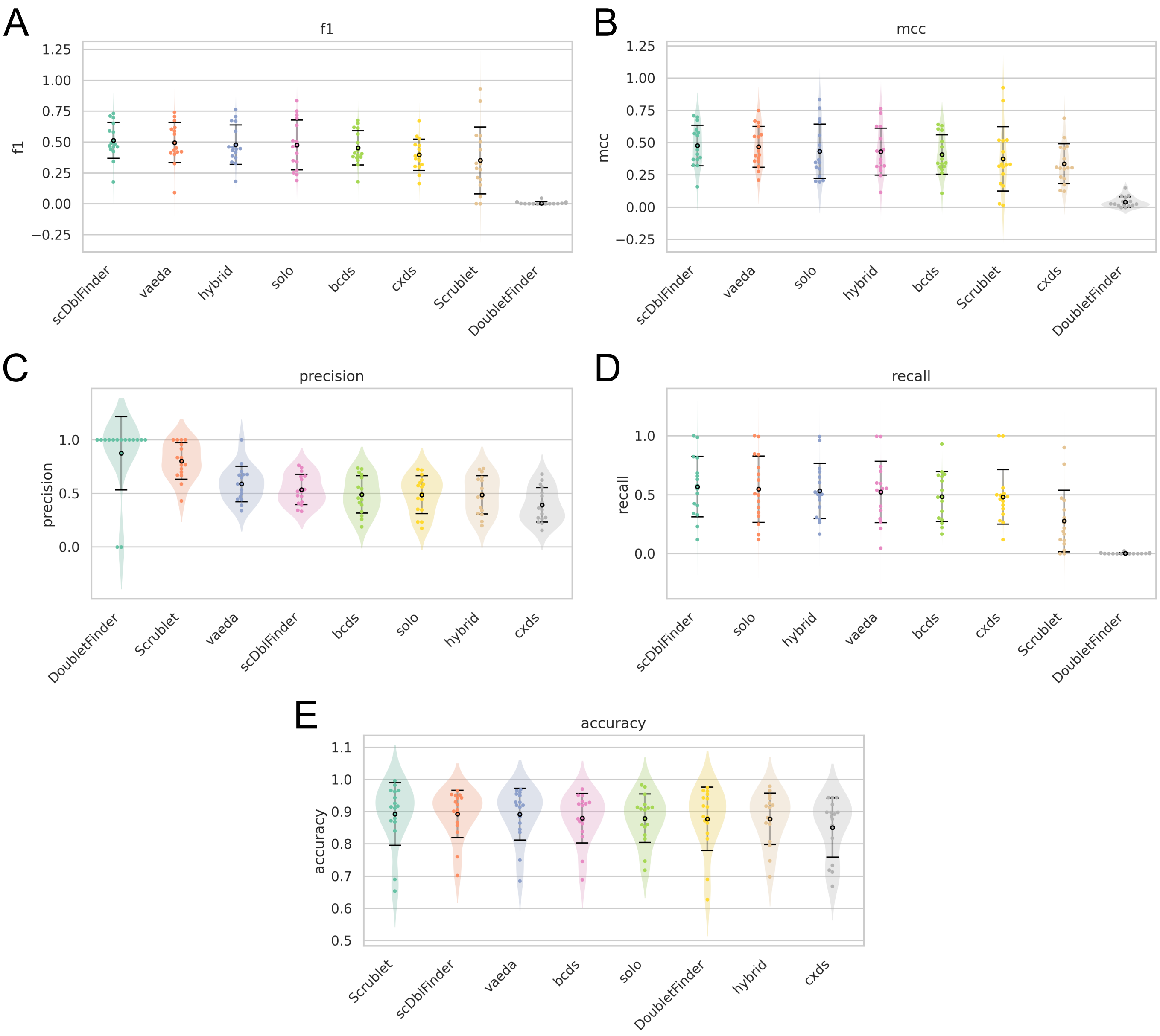


Supplemental Figure S8: ***Doublet Calling performance varies highly with dataset.*** Shown are violin plots of metrics (f1 (A), mcc(B), precision(C), recall(D), and accuracy (E)) for accessing doublet calling. Methods are given on the x axis and ordered by decreasing performance. Each point represents one of the benchmarking datasets and the black point is the mean. Error bars represent the variance.

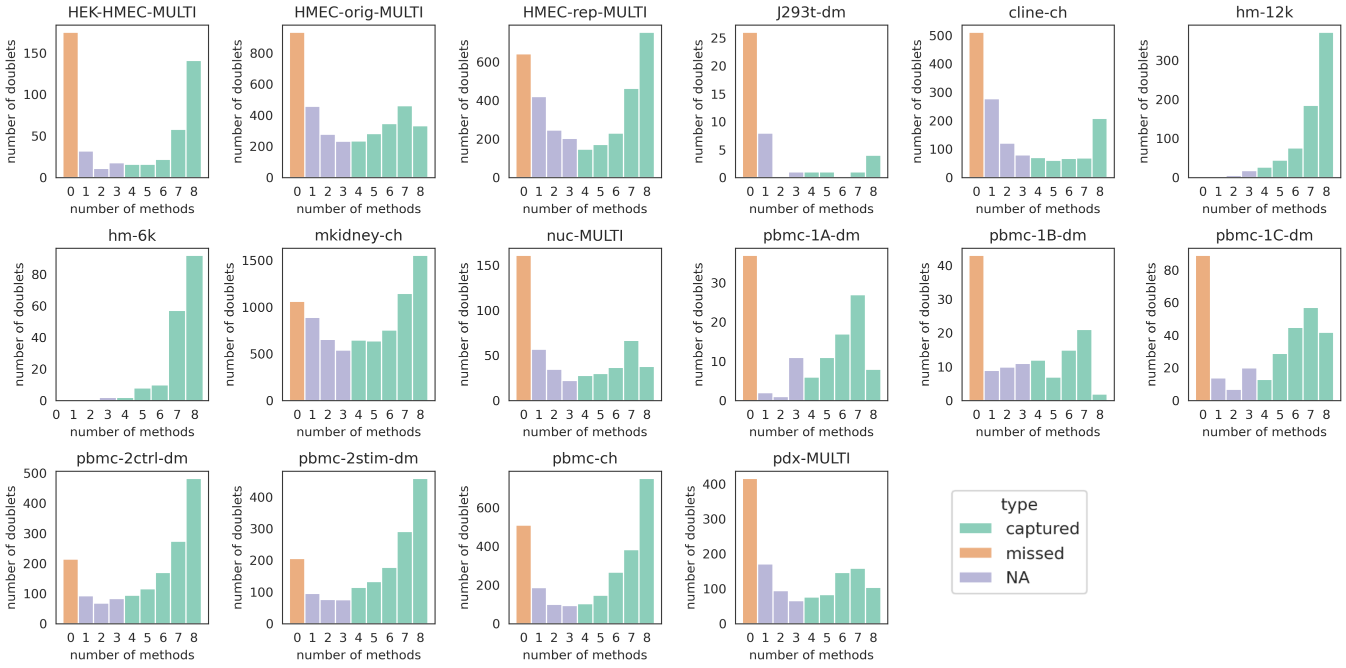


Supplemental Figure S9: ***Most datasets have a sizable population of doublets that are misclassified by every method.*** Depicted above are histograms of the number of doublets that are classified by 0, 1, 2, 3, 4, 5, 6, or 8 of the methods. We define ”missed” doublets as doublets that are classified by none of the methods and ”captured” doublets as doublets that are classified by at least 4 of the methods.

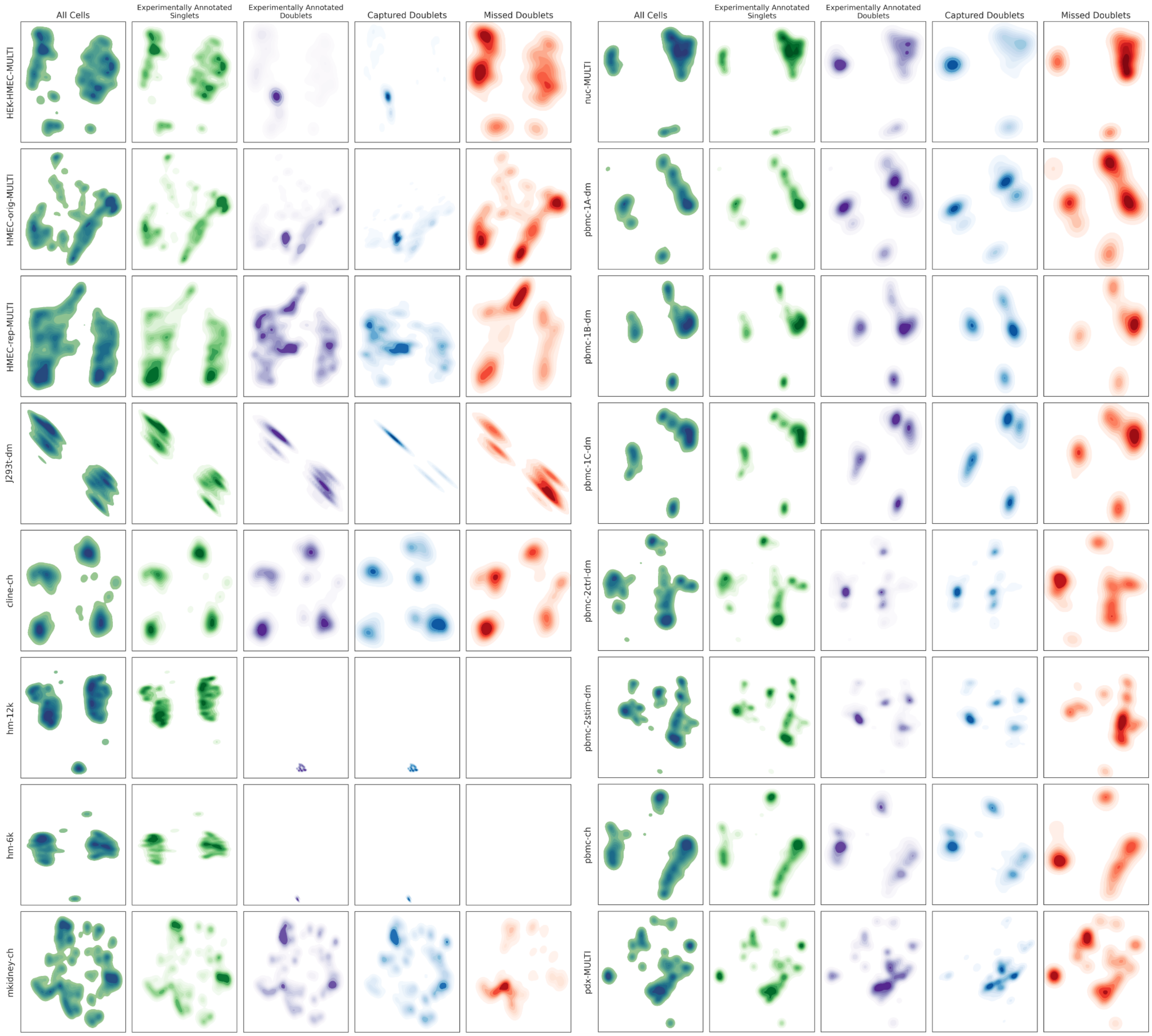


Supplemental Figure S10: ***Doublets misclassified by all methods have greater mixing with singlets.*** Shown are UMAP projections of top 30 principal components of each dataset. Rows are the benchmarking datasets. The first column in each panel are all cells in the dataset; the second and third columns show experimentally annotated singlets and doublets, respectively; the third column shoes "captured" doublets, and the fourth column shows "missed" doublets. Where "captured" doublets are experimentally annotated doublets classified by at least 4 of the 8 methods and "missed" doublets are experimentally annotated doublets that are misclassified by all of the methods.

### S2 Supplemental Tables

| dataset | number of cells | number of doublets | doublet fraction | median UMI | median gene count |
|---|---|---|---|---|---|
| HEK-HMEC-MULTI | 10,641 | 489 | 0.046 | 17,424 | 3,795 |
| HMEC-orig-MULTI | 26,426 | 3,568 | 0.135 | 23,502 | 4,598 |
| HMEC-rep-MULTI | 10,580 | 3,282 | 0.310 | 1,188 | 601 |
| J293t-dm | 500 | 42 | 0.084 | 14,135 | 3,461 |
| cline-ch | 7,954 | 1,465 | 0.184 | 4,825 | 2,149 |
| hm-12k | 12,820 | 730 | 0.057 | 12,424 | 3,147 |
| hm-6k | 6,806 | 171 | 0.025 | 21,301 | 4,032 |
| mkidney-ch | 21,179 | 7,901 | 0.373 | 3,929 | 1,687 |
| nuc-MULTI | 5,578 | 475 | 0.085 | 1,021 | 786 |
| pbmc-1A-dm | 3,298 | 120 | 0.036 | 973 | 384 |
| pbmc-1B-dm | 3,790 | 130 | 0.034 | 862 | 361 |
| pbmc-1C-dm | 5,270 | 316 | 0.060 | 829 | 352 |
| pbmc-2ctrl-dm | 13,913 | 1,598 | 0.115 | 1,276 | 526 |
| pbmc-2stim-dm | 13,916 | 1,631 | 0.117 | 1,361 | 550 |
| pbmc-ch | 15,272 | 2,545 | 0.167 | 556 | 323 |
| pdx-MULTI | 10,296 | 1,317 | 0.128 | 2,243 | 1,029 |

Supplemental Table S1: Benchmarking datasets.

| | vaeda-scDblFinder | vaeda-DoubletFinder | scDblFinder-DoubletFinder |
|---|---|---|---|
| HEK-HMEC-MULTI | vaeda | vaeda | ≈ |
| HMEC-orig-MULTI | vaeda | vaeda | scDblFinder |
| HMEC-rep-MULTI | vaeda | ≈ | DoubletFinder |
| J293t-dm | ≈ | DoubletFinder | DoubletFinder |
| cline-ch | scDblFinder | ≈ | scDblFinder |
| hm-12k | ≈ | DoubletFinder | DoubletFinder |
| hm-6k | ≈ | DoubletFinder | DoubletFinder |
| mkidney-ch | scDblFinder | vaeda | scDblFinder |
| nuc-MULTI | scDblFinder | DoubletFinder | ≈ |
| pbmc-1A-dm | scDblFinder | vaeda | scDblFinder |
| pbmc-1B-dm | scDblFinder | vaeda | scDblFinder |
| pbmc-1C-dm | scDblFinder | DoubletFinder | scDblFinder |
| pbmc-2ctrl-dm | scDblFinder | ≈ | scDblFinder |
| pbmc-2stim-dm | scDblFinder | ≈ | scDblFinder |
| pbmc-ch | scDblFinder | vaeda | scDblFinder |
| pdx-MULTI | scDblFinder | vaeda | scDblFinder |

Supplemental Table S2: *Comparison of top three methods stratified by dataset.* Wilcoxon rank-sum tests were used to identify significant performance (p¡=0.05) differences between the top three methods on the 95% down-samplings. ≈ implies $p > 0.05$.

| | f1 | mcc | precision | recall | accuracy |
|---|---|---|---|---|---|
| scDblFinder | 50.1 | 46.9 | 55.5 | 55.6 | 88.8 |
| vaeda | 48.5 | 45.3 | 56.4 | 53.9 | 88.3 |
| hybrid | 45.9 | 42.6 | 53.8 | 51.4 | 88.0 |
| solo | 44.9 | 41.1 | 49.9 | 50.3 | 88.0 |
| bcds | 45.3 | 41.4 | 50.4 | 50.5 | 87.9 |
| cxds | 39.0 | 34.6 | 46.2 | 44.7 | 86.5 |
| Scrublet | 38.8 | 34.4 | 45.9 | 44.3 | 86.5 |
| lib-sze | 27.5 | 20.6 | 29.5 | 30.7 | 84.5 |
| DoubletFinder | 45.1 | 41.3 | 52.9 | 50.3 | 87.5 |

Supplemental Table S3: Doublet classification across 16 benchmark datasets (averaged performance). Instead of using each method's doublet callers, we took the top $n^2 * 10^{-5}$ scoring doublets.

| | f1 | mcc | precision | recall | accuracy |
|---|---|---|---|---|---|
| scDblFinder | 1.3 | 0.9 | -1.8 | 1.3 | 0.5 |
| vaeda | 1.2 | 1.4 | 2.5 | -1.4 | 0.9 |
| hybrid | 1.9 | 0.4 | -5.0 | 1.9 | -0.3 |
| solo | 2.7 | 2.2 | -1.1 | 4.5 | -0.1 |
| bcds | -0.1 | -0.5 | -1.4 | -1.9 | 0.1 |
| cxds | 0.6 | -0.9 | -6.9 | 3.6 | -1.4 |
| Scrublet | -3.6 | 3.1 | 34.4 | -16.6 | 2.8 |
| lib-sze | 0.0 | 0.0 | 0.0 | 0.0 | 0.0 |
| DoubletFinder | -44.4 | -37.2 | 0.346 | -50.0 | 0.3 |

Supplemental Table S4: Difference between doublet caller metrics and expected doublet classification metrics (negative means using expected number of doublets performs better than the methods built in doublet caller)
